# Type I and III interferons differentially encode inflammatory gene activation through threshold control of ISGF3 dynamics

**DOI:** 10.64898/2026.10.01.756040

**Authors:** Henrry Dubon Martinez, Rajshree Chettiyar, Maya McAuley, Catera L. Wilder

## Abstract

Type I and type III interferons (IFNs) activate the shared Janus kinase/signal transducer and activator of transcription (JAK-STAT) pathway but generate distinct inflammatory transcriptional responses in addition to their shared antiviral transcriptional induction. While some mechanisms including cytokine-receptor affinity and receptor abundance have been proposed for regulating type I vs type III IFN specificity, they account for differences in signal amplitude but do not fully explain IFN-specific changes in JAK-STAT activity over time (i.e. the signaling temporal dynamics) which has been shown to be predictive of inflammatory gene activation. Here, we investigated mechanisms that could explain how type I and III IFNs produce distinct temporal dynamics of the IFN-activated transcription factor IFN-stimulated gene factor 3 (ISGF3), leading to IFN-type specific activation of inflammatory genes in lung epithelial cells. We show that IFN-β, a type I IFN, and IFN-λ3, a type III IFN, induced comparable initial ISGF3 activation but subsequently generated distinct signaling trajectories, with IFN-β undergoing potentiation and IFN-λ3 limiting the duration of high ISGF3 activity, preventing robust inflammatory gene induction despite prolonged lower-level activity. Inflammatory gene induction was associated with the magnitude and duration of ISGF3 activity above an activation threshold during the intermediate phase of the signal response. The characteristic IFN-λ3-induced ISGF3 signaling dynamics were maintained at saturating doses of IFN-λ3 indicating the lack of signal activation above the threshold was independent of the dose. Additionally, differences in the abundance of the ISGF3 complex subunits STAT1, STAT2, and interferon regulatory factor 9 (IRF9), or in the induction of known negative regulators did not readily explain the response. Blocking protein synthesis caused rapid loss of IFN-λ3-induced STAT phosphorylation and ISGF3 activity, supporting the presence of a pre-existing negative regulator limiting IFN-λ3-mediated JAK-STAT pathway activation above the threshold required for activating inflammatory genes, while IFN-β and IFN-λ3 co-stimulation localized this regulation upstream of the shared ISGF3 complex. Increasing pathway competence through IFN-β priming enhanced the IFN-λ3-induced ISGF3 response above the inflammatory gene activation threshold leading to induction of inflammatory genes, albeit transiently. Together, these findings support a model in which the magnitude and duration of ISGF3 activity above an inflammatory gene activation threshold, rather than pathway activation alone, contribute to stimulus-specific transcriptional responses and suggest that a negative regulatory mechanism upstream of the ISGF3 complex constrains the temporal dynamics of IFN-λ3 signaling.

## INTRODUCTION

Interferons (IFNs) are central mediators of innate antiviral immunity, coordinating cellular responses to viral infection (1). Although type I and III interferons (IFNs) activate the shared Janus kinase/signal transducers and activators of transcription (JAK-STAT) pathway and antiviral programs, their protective versus potentially pathogenic immune functions are controlled by markedly different induction of inflammatory responses (2). These differences are not explained by activation of the core JAK-STAT pathway, as both type I and III IFNs induce STAT1 and STAT2 phosphorylation and their association with interferon regulatory factor 9 (IRF9) to form interferon-stimulated gene factor 3 (ISGF3), which translocates to the nucleus and induces interferon-stimulated genes (ISGs) (3, 4). Prior work has proposed receptor abundance as one determinant of these differences. In hepatocytes, low abundance of the type III IFN receptor, IFNλ receptor (IFNLR) can limit STAT1 activation and IRF1 induction, thereby restricting an IRF1-dependent inflammatory program (5). In contrast, studies in intestinal epithelial cells found that increasing receptor abundance altered the amplitude of the response but did not account for the characteristic temporal dynamic differences between type I and type III IFNs (6). Together, these findings suggest that receptor abundance can influence signaling capacity, but additional mechanisms may determine how the shared JAK–STAT pathway is regulated over time.

Temporal control is an important source of signaling specificity in pathways that use shared molecular components (7). We previously showed that IFN-β, a type I IFN, induces a second, potentiated phase of ISGF3 activity that depends on stimulus dose, duration, and positive feedback through STAT2 and IRF9 (8). Conditions that fail to engage this potentiated phase, including low-dose or short-duration stimulation with IFN-β and stimulation with IFN-λ3, a type III IFN, also fail to robustly induce a subset of inflammatory genes. These observations raised a mechanistic question that could not be resolved by simply measuring whether ISGF3 or IRF1 was on or off: which features of the signaling trajectory determine whether inflammatory genes are activated by an IFN response, and what mechanisms regulate these features?

Here, we investigate mechanisms that control the temporal regulation of ISGF3 activity and distinguish IFN-β and IFN-λ3 responses in lung epithelial cells. Using quantitative, time-resolved measurements and targeted perturbations, we examine the mechanisms underlying the distinct temporal dynamics of IFN-λ3-mediated JAK-STAT signaling. We find that inflammatory gene induction is associated with ISGF3 activity exceeding an activation threshold during a critical intermediate phase of the response during which IFN-3-mediated signaling experiences a transient downregulation followed by pathway reactivation which we have termed a trough. Although IFN-λ3 can induce IRF1, IRF1 expression is rapidly truncated, and saturating doses of IFN-λ3 does not eliminate the characteristic ISGF3 trough. The trough is also not explained by insufficient abundance of STAT1, STAT2, or IRF9, or by preferential induction of SOCS1, SOCS3, or USP18. Instead, protein-synthesis inhibition reveals a negative regulatory activity independent of protein synthesis associated with IFN-λ3-mediated JAK-STAT signaling, and co-stimulation places this regulation upstream of the shared ISGF3 complex. Finally, IFN-β priming increases the initial IFN-λ3-induced ISGF3 response sufficiently to cross the inflammatory gene activation threshold and transiently induce inflammatory genes despite reduced IRF1 induction. Together, these findings suggest that mechanisms controlling the temporal dynamics of ISGF3 activity determine whether JAK-STAT signaling crosses gene-specific activation thresholds and engages inflammatory transcriptional programs.

## RESULTS

### Differences in ISGF3 temporal dynamics are sufficient to cause type I and III IFN-specific induction of inflammatory genes

To examine how type I and type III IFNs generate distinct signaling dynamics through the shared JAK–STAT pathway (Fig. 1A), we stimulated the murine MLE-12 lung epithelial cells with either 10 U/ml IFN-β or 100 ng/ml IFN-λ3 and collected cytoplasmic and nuclear extracts at various time points post stimulation. These concentrations were selected so that the initial activation of the IFN signaling pathway was comparable between IFN-β and IFN-λ3 as measured by ISGF3 activity at 30 min. We measured nuclear ISGF3 activity using an electrophoretic mobility shift assay (EMSA) with an ISRE oligo probe using constitutive transcription factor NFY as control, while the subunits of ISGF3, phosphorylated STAT1 (pSTAT1), phosphorylated STAT2 (pSTAT2), and IRF9, that were in the cytoplasm were measured using immunoblotting (Fig. 1B). With the IFN-λ3 stimulation, after an initial robust activation during the first hour, nuclear ISGF3 and cytoplasmic pSTAT1 and pSTAT2 activation decreased starting around one hour, with ISGF3 activity decreasing almost 40% compared to the activity during the initial activation period (Fig. 1C). This inactivation was transient in nature as nuclear ISGF3 and cytoplasmic pSTAT1 and pSTAT2 began increasing again after 2 hours post stimulation. This transient downregulation of ISGF3 1 to 2 hours post stimulation was specific to IFN-λ3 stimulation and not IFN-β.

**Figure 1.**
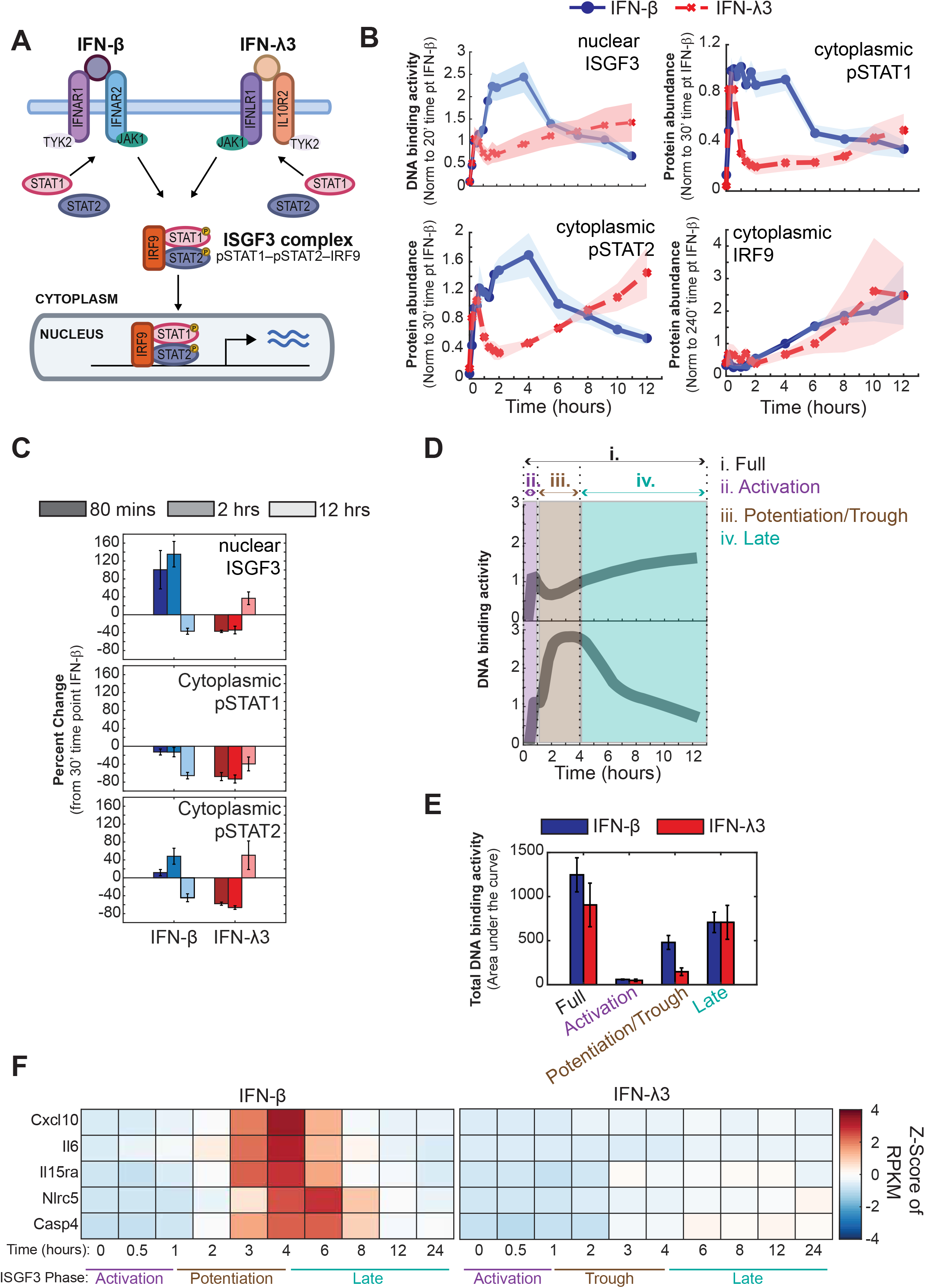
Induction of pro-inflammatory genes by type I IFN but not type III correlates with distinct temporal dynamics of ISGF3 activation. A. Schematic of type I and III IFN signaling through the JAK-STAT pathway and downstream formation of the ISGF3 complex. B. Time course of ISGF3 DNA-binding activity and cytoplasmic abundance of ISGF3 components following stimulation with 10 U/mL IFN-β (solid blue) or 100 ng/mL IFN-λ3 (dashed red). Nuclear ISGF3 DNA-binding activity was measured by EMSA, and cytoplasmic pSTAT1, pSTAT2, and IRF9 abundance was quantified by western blotting. Data were normalized to IFN-β stimulation at 20 min for ISGF3, 30 min for pSTAT1 and pSTAT2, and 240 min for IRF9. Shaded areas represent SEM from at least five independent experiments (n = 5). C. Bar graphs depicting percent change in nuclear ISGF3 DNA-binding activity, and cytoplasmic pSTAT1 and pSTAT2 abundance at 80 min, 2 h, and 12 h following IFN-β and IFN-λ3 stimulation. Values were calculated relative to the response at 30 min of IFN-β stimulation. Error bars represent SEM from at least three independent experiments (n = 3). D. Schematic of area under the curve (AUC) to quantify ISGF3 DNA-binding activity over the full response (i), the activation (ii), potentiation/trough (iii), and late (iv) phases. E. ISGF3 DNA-binding activity quantified by AUC following stimulation with 10 U/ml IFN-β (blue) or 100 ng/ml IFN-λ3 (red) over the full-time course, activation, potentiation, and late phases. Bars represent ± SEM from 3 independent experiments (n = 3). F. Heatmap of RNA-seq expression profiles of selected IFN-responsive genes following stimulation with 10 U/mL IFN-β or 100 ng/mL IFN-λ3 over the indicated time course. Mean RPKM values from two biological replicates were Z-score normalized for each gene. Red and blue indicate expression above and below the gene-specific mean, respectively. ISGF3 signaling phases defined in (D) are indicated below the heatmap.

To determine if features of the ISGF3 temporal trajectories were important for inducing inflammatory gene expression, we identified three phases of the temporal dynamics of ISGF3 when stimulated with IFN-β or -λ3: activation phase, potentiation/trough phase, and late phase (Fig. 1D). We determined the total ISGF3 activity by calculating the area under the curve (i.e. the integral) during the entire time course or for each phase and found that of the three different phases of the ISGF3 temporal trajectories, the potentiation/trough phase had the greatest difference in ISGF3 activity between IFN-β and -λ3 (Fig. 1E). To assess if the difference in total ISGF3 activity by IFN-β and -λ3 during the potentiation/trough phase had any functional consequences, we plotted gene expression of pro-inflammatory genes from our previously published RNA sequencing (RNAseq) dataset (8). Pro-inflammatory gene induction by IFN-β was initiated between 2-3 hours which correlated with robust ISGF3 activity by IFN-β during the potentiated phase (FDR corrected p-value < 0.05) (Fig. 1F). This was in contrast to no significant induction of pro-inflammatory gene expression by IFN-3, correlating with diminished ISGF3 activity during the trough phase.

### Activation thresholds of ISGF3 control stimulus-specific gene expression through shared JAK-STAT pathway

Since previous work demonstrated that differential activation of the transcription factor IRF1 contributes to the distinct inflammatory gene expression responses induced by type I and type III IFNs, we asked whether differential IRF1 induction may similarly contribute to the distinct inflammatory responses induced by IFN-β and IFN-λ3 in lung epithelial cells (5). To address this question, we examined the mRNA expression of IRF1 from the RNAseq dataset of MLE-12 lung epithelial cells stimulated with IFN-β or IFN-λ3 and mRNA collected at various time points post stimulation (Fig. 2A). Both IFN-β and IFN-λ3 induced IRF1 expression within the first hour of stimulation (FDR corrected p-value < 0.05). After an hour, IRF1 expression continued to increase with IFN-β stimulation, peaking at approximately two hours before beginning to decline after four hours. In contrast, IFN-λ3-induced IRF1 expression was rapidly truncated after the first hour and subsequently maintained at lower levels. These results indicate that IFN-λ3 can initiate IRF1 expression in lung epithelial cells but cannot sustain its induction to the same extent as IFN-β.

**Figure 2.**
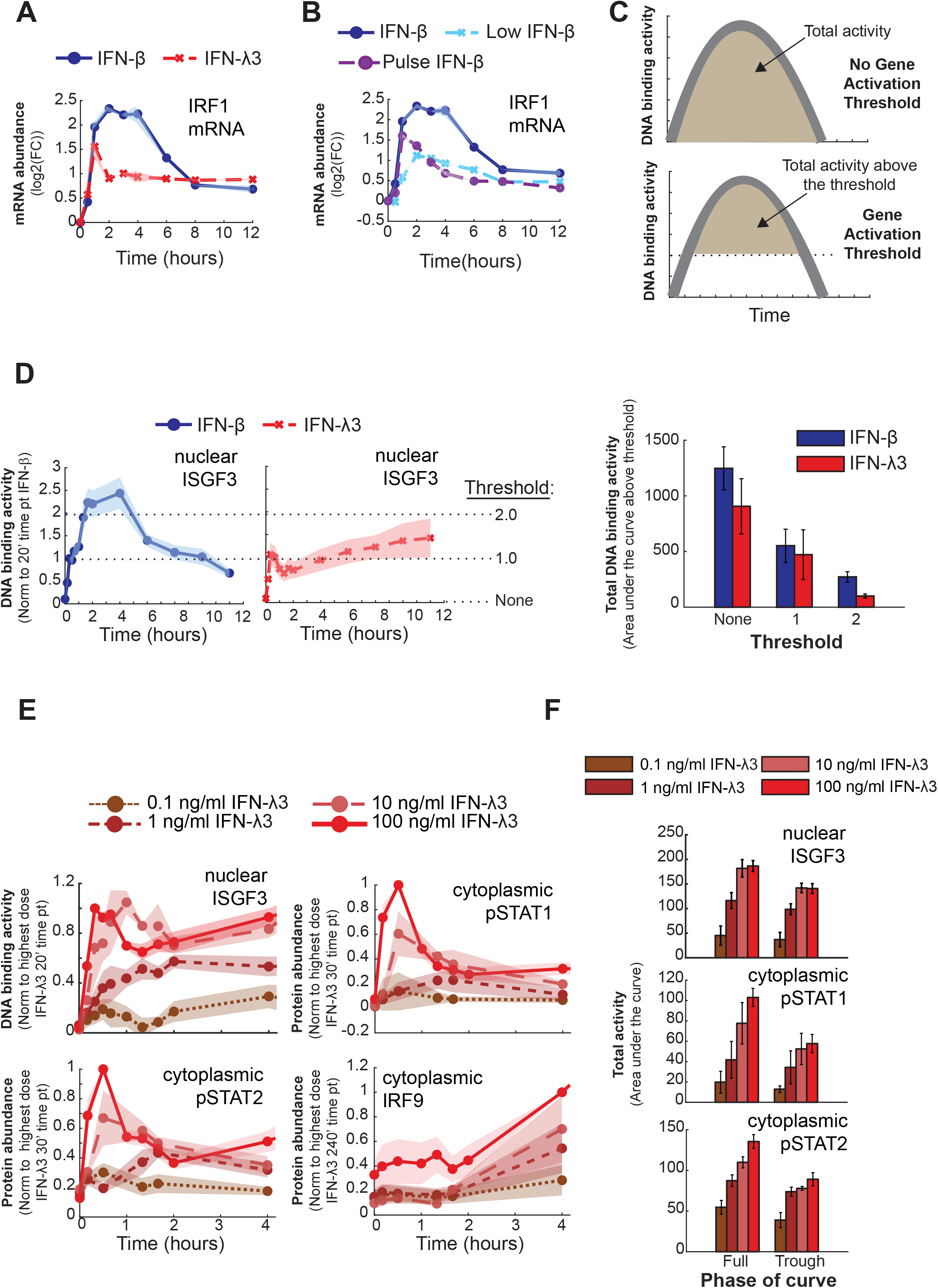
High doses of IFNλ3 do not alter the signaling dynamics to potentiate above the activation threshold. A. Temporal dynamics of IRF1 mRNA expression following stimulation with 10 U/mL IFN-β (solid blue) or 100 ng/mL IFN-λ3 (dashed red). IRF1 mRNA abundance was measured by RNAseq, and shown as log₂ fold change. Shaded areas represent SEM from two independent experiments (n = 2). B. IRF1 mRNA expression following stimulation with IFN-b (10 U/mL, solid blue), low IFN-b (1 U/mL, dashed light blue), or a 15 min pulse of 10 U/mL IFN-b (dashed purple). IRF1 mRNA abundance was measured as described in (A). Shaded areas represent SEM from two independent experiments (n = 2). C. Schematic illustrating AUC analysis of ISGF3 DNA-binding activity in the absence or presence of a gene-activation threshold. D. ISGF3 DNA-binding activity following stimulation with 10 U/mL IFN-b (solid blue) or 100 ng/mL IFN-λ3 (dashed red), illustrating the thresholds defined in (C). AUC was calculated above the indicated DNA-binding activity thresholds (none, 0.5, 1.0) and quantified for IFN-b and IFN-λ3. Bars represent mean ± SEM from three independent experiments (n = 3). E. Nuclear ISGF3 DNA-binding activity and cytoplasmic abundance of ISGF3 components following stimulation with increasing concentrations of IFN-l3 (0.1 - 100 ng/ml). ISGF3 DNA-binding activity was measured by EMSA and normalized to 100 ng/mL IFN-l3 at 20 min. Cytoplasmic abundance of the pSTAT1, pSTAT2, and IRF9 was measured by western blotting and normalized to 100 ng/mL IFN-l3 stimulation at 30 min for pSTAT1 and pSTAT2 and 240 min for IRF9. Shaded areas depict the mean ± SEM from at least 3 independent experiments (n = 3). F. AUC quantification of nuclear ISGF3 DNA-binding activity and cytoplasmic pSTAT1 and pSTAT2 abundance following stimulation with increasing concentrations of IFN-λ3 (0.1 - 100 ng/mL), as shown in (E). AUC was calculated over the full-time course or during the trough phase (80 min). Bars represent mean ± SEM from three independent experiments (n = 3).

We next asked whether truncated IRF1 induction was specific to IFN-λ3 or more generally associated with IFN stimulation conditions that fail to induce the inflammatory gene expression program. We previously reported that either reducing the dose of IFN-β or limiting IFN-β exposure to a short pulse diminished the potentiated phase of ISGF3 activity and prevented robust induction of inflammatory genes (8). We therefore examined IRF1 expression when MLE-12 lung epithelial cells were stimulated with low IFN-β (1 U/ml) or pulse IFN-β (15 minutes of 10 U/ml), from the same RNAseq dataset. Like IFN-λ3, both low IFN-β and pulse IFN-β stimulation reduced IRF1 expression after an initial induction compared with the sustained IRF1 response seen with IFN-β stimulation (Fig. 2B). Thus, reduced IRF1 induction was associated with multiple stimulation conditions that failed to generate the potentiated ISGF3 response and the inflammatory gene expression program.

These observations led us to consider whether inflammatory gene induction may require ISGF3 activity to exceed a critical activation threshold (Fig. 2C). Rather than considering only the total amount of ISGF3 activity generated over time, we reanalyzed the IFN-β- and IFN-λ3-induced ISGF3 signaling trajectories by quantifying the integrated ISGF3 activity with no threshold or with thresholds such that only ISGF3 DNA-binding activity above the threshold was used to determine the area under the curve (Fig. 2D). When no activation threshold was imposed, IFN-β and IFN-λ3 generated more comparable amounts of integrated ISGF3 activity. However, as the activation threshold was increased, progressively more of the IFN-λ3 response fell below the threshold, whereas IFN-β maintained substantial ISGF3 activity above it. These results suggest that the distinction between IFN-β and IFN-λ3 induction of pro-inflammatory genes may not be determined simply by total ISGF3 activity, but by the amount of ISGF3 activity that exceeds a threshold required for induction of more stringently regulated genes, including inflammatory genes.

We next asked whether the inability of IFN-λ3 to sustain ISGF3 activity above this threshold was due to the dose of IFN-λ3 being used. To address this, MLE-12 cells were stimulated with increasing concentrations of IFN-λ3 ranging from 0.1 ng/ml to 100 ng/ml, and nuclear ISGF3 DNA-binding activity and the cytoplasmic abundance of phosphorylated STAT1 and STAT2 were measured over time using EMSA and immunoblots, respectively (Fig. 2E). Increasing the IFN-λ3 dose increased pathway activation at lower concentrations. However, the ISGF3 response approached saturation at the two highest doses tested, 10 and 100 ng/ml IFN-λ3. We measured the integrated ISGF3 activity in addition to the total abundance of cytoplasmic pSTAT1 and pSTAT2 for the entire time course and for the trough phase and found that increasing the dose from 10 to 100 ng/ml produced little additional increase in either the activation of ISGF3 during the entire time course or during the trough phase (Fig. 2F). Similar saturation was observed in the upstream abundance of pSTAT1 and pSTAT2. Thus, high IFN-λ3 concentrations are insufficient to convert the characteristic IFN-λ3 signaling trajectory into the sustained, high-amplitude response observed with IFN-β.

### Positive feedback regulator abundance does not account for the attenuated IFN-λ3 signaling trajectory

Having determined that the dose of IFN-λ3 was insufficient to increase ISGF3 activity above the threshold associated with inflammatory gene induction, we next investigated mechanisms that may constrain the IFN-λ3 signaling response. We previously reported that the potentiated phase of ISGF3 activity induced by IFN-β depends on a positive feedback loop involving the ISGF3 components STAT2 and IRF9 (8). Disrupting the inducibility of these components diminished the potentiated phase of ISGF3 activity and reduced inflammatory gene expression. We therefore asked whether the inability of IFN-λ3 to generate ISGF3 activity above the inflammatory activation threshold could result from insufficient induction of components of this positive feedback loop.

To address this question, we measured the temporal expression of STAT1, STAT2, and IRF9 mRNA following stimulation of MLE-12 lung epithelial cells with IFN-β or IFN-λ3 (Fig. 3A). Both IFNs induced expression of STAT1, STAT2, and IRF9 with generally similar temporal profiles. Although differences in STAT1 and STAT2 mRNA abundance were observed at later time points between four and eight hours, these differences occurred after the period in which the IFN-β and IFN-λ3 ISGF3 trajectories had already diverged.

**Figure 3.**
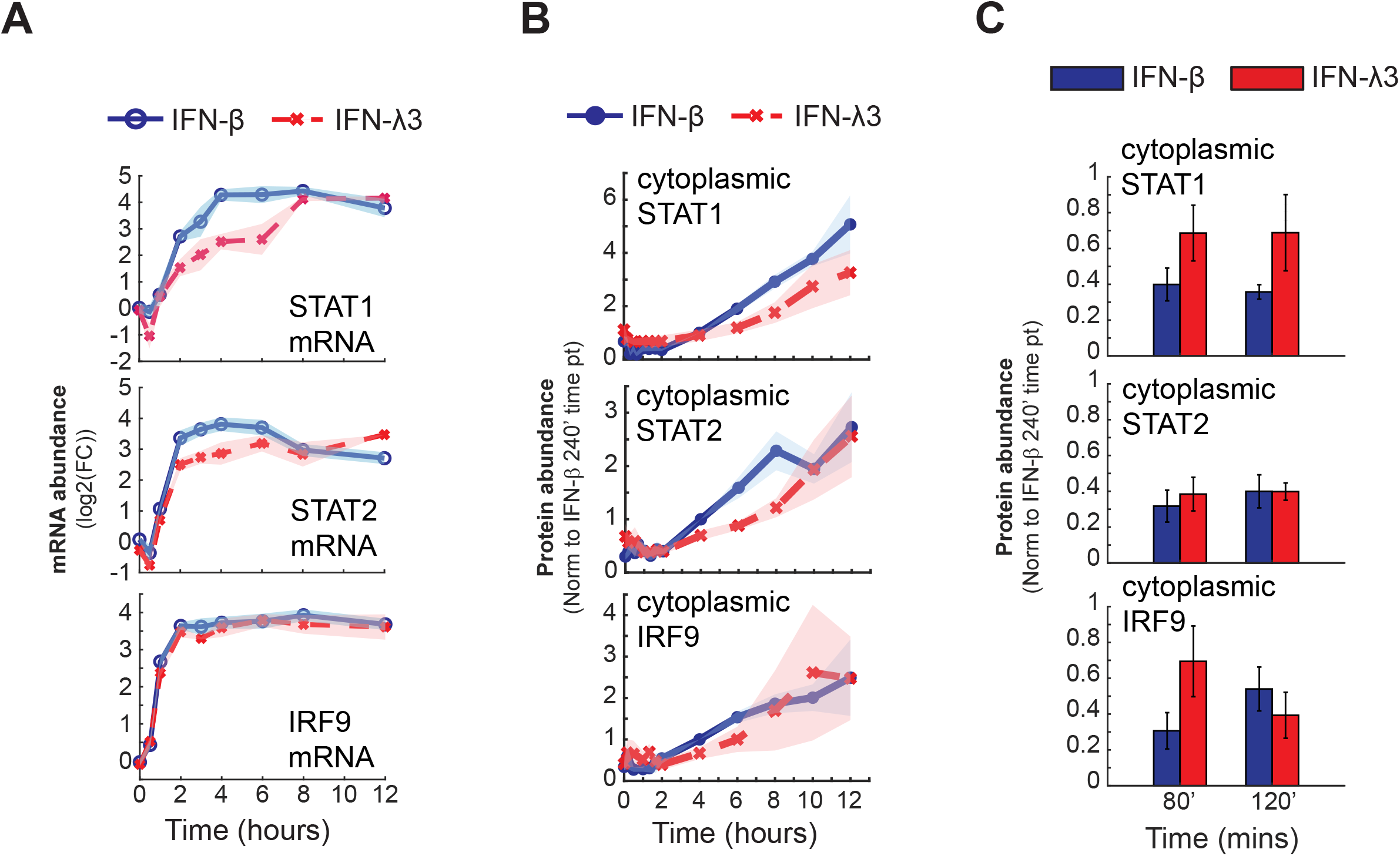
Differences in type I and III IFN temporal dynamic control are not due to IFN-type specific induction of positive regulators STAT1, STAT2, and IRF9. A. STAT1, STAT2, and IRF9 mRNA expression following stimulation with 10 U/mL IFN-β (solid blue) or 100 ng/mL IFN-λ3 (dashed red). RT-qPCR analysis and normalization were performed as described in Figure 2A. Shaded areas represent SEM from three independent experiments (n = 3). B. Cytoplasmic STAT1, STAT2, and IRF9 abundance following stimulation as described in (B). Protein abundance was quantified by western blotting and normalized to 10 U/mL IFN-β stimulation at 240 min. Shaded areas represent SEM from three independent experiments (n = 3). C. Cytoplasmic STAT1, STAT2, and IRF9 abundance at 80 min, corresponding to the trough phase, and 120 min following IFN-β or IFN-λ3 stimulation. Protein abundance was quantified and normalized as described in (C). Bars represent mean ± SEM from three independent experiments (n = 3).

We next determined whether differences in mRNA induction resulted in differences in the abundance of the corresponding signaling proteins. Cytoplasmic STAT1, STAT2, and IRF9 protein abundance was measured over a 12-hour time course following IFN-β or IFN-λ3 stimulation (Fig. 3B). Despite some differences in mRNA expression, the abundance of the three proteins was generally similar between the two stimulation conditions. We specifically examined the 80 mins and two-hour time points since they correspond to the potentiated phase of IFN-β-induced ISGF3 activity and the trough phase of the IFN-λ3 response (Fig. 3C). During this interval, IFN-λ3-stimulated cells did not have a lower abundance of STAT1, STAT2, or IRF9 and, at some time points, showed similar or greater amounts of these proteins than IFN-β-stimulated cells.

Together, these results indicate that the attenuated ISGF3 activity observed during the IFN-λ3 trough phase cannot be explained by insufficient abundance of the STAT1, STAT2, or IRF9 components required to form ISGF3. Thus, although positive feedback regulation of STAT2 and IRF9 contributes to the potentiated ISGF3 response induced by IFN-β, differences in the abundance of these positive regulators do not appear sufficient to explain why IFN-λ3 fails to sustain ISGF3 activity above the threshold associated with inflammatory gene induction.

### The IFN-λ3 signaling trough is associated with transient inactivation upstream of nuclear ISGF3

Having determined that the reduced ISGF3 activity during the IFN-λ3 trough phase was not caused by insufficient abundance of the positive regulators STAT1, STAT2, or IRF9, we next asked where within the signaling pathway the IFN-λ3-specific inactivation occurred. Because ISGF3 activity was measured in nuclear extracts, one possibility was that the trough reflected differences in retention of activated STAT proteins in the cytoplasm rather than in the inactivation of signaling upstream of ISGF3 formation. We therefore compared the temporal abundance of phosphorylated and total STAT1 and STAT2 in the nuclear and cytoplasmic fractions of MLE-12 cells stimulated with IFN-β or IFN-λ3 (Fig. 4A).

**Figure 4.**
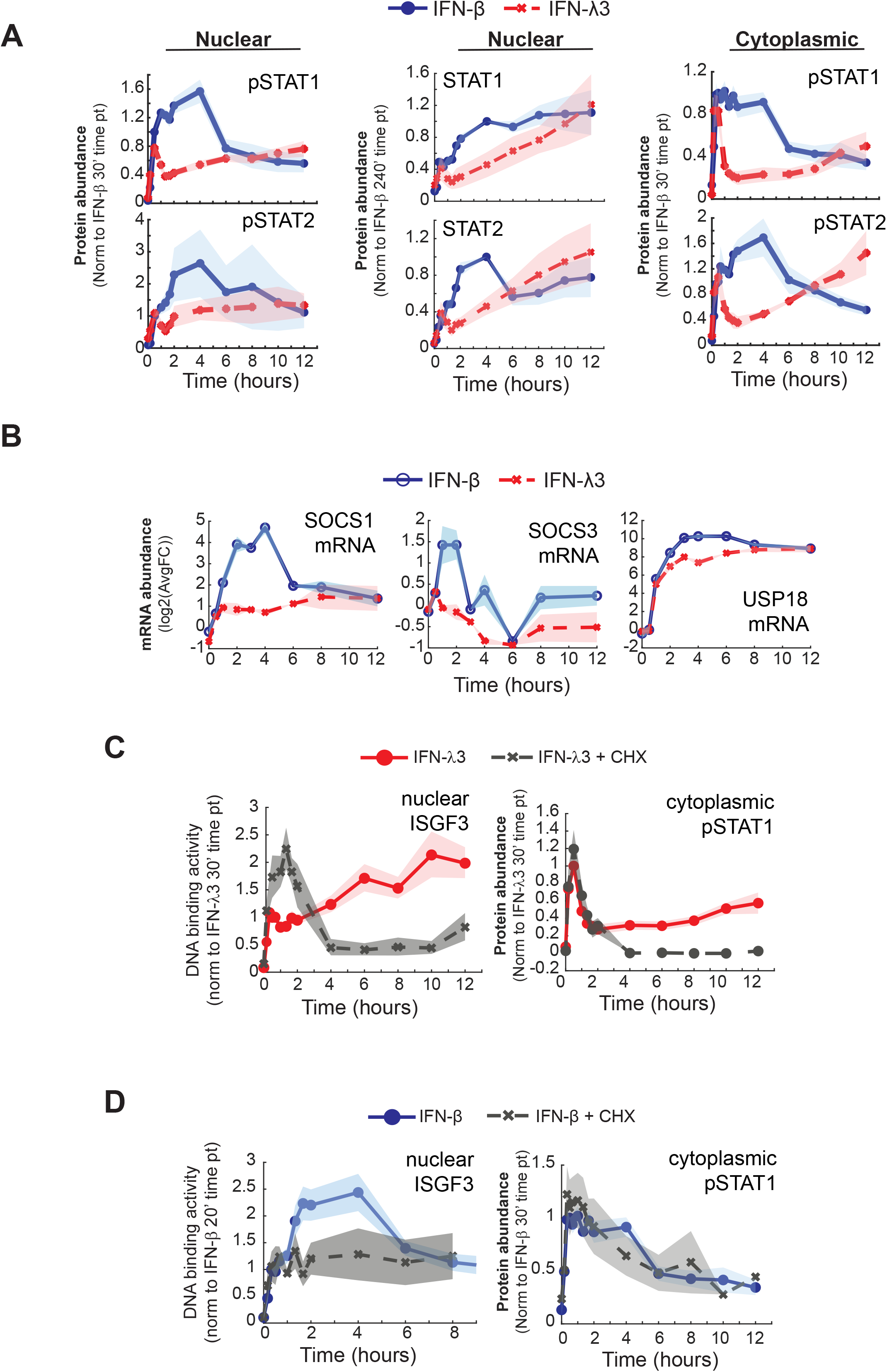
The stability of the pathway activation is different between type I and III IFNs, indicating differences in pathway negative regulators. A. Temporal dynamics of nuclear pSTAT1, pSTAT2, STAT1, and STAT2 and cytoplasmic pSTAT1 and pSTAT2 abundance following stimulation with 10 U/mL IFN-β (solid blue) or 100 ng/mL IFN-λ3 (dashed red). Protein abundance was quantified by western blotting and normalized to IFN-β stimulation at 30 min for pSTAT1 and pSTAT2 and 240 min for STAT1 and STAT2. Shaded areas represent SEM from at least three independent experiments (n = 3). B. SOCS1, SOCS3, and USP18 mRNA expression following stimulation with 10 U/mL IFN-β (solid blue) or 100 ng/mL IFN-l3 (dashed red). mRNA quantification and normalization were performed as described in Figure 2A. Shaded areas represent SEM from two independent experiments (n = 2). C. Nuclear ISGF3 DNA-binding activity and cytoplasmic pSTAT1 abundance following stimulation with 100 ng/mL IFN-λ3 in the presence (gray) or absence (red) of 20 µg/mL cycloheximide (CHX. ISGF3 DNA-binding activity and pSTAT1 abundance were measured by EMSA and western blotting, respectively, and normalized to IFN-λ3 stimulation at 30 min. Shaded areas represent SEM from at least three independent experiments (n = 3). D. Nuclear ISGF3 DNA-binding activity and cytoplasmic pSTAT1 abundance following 10 U/mL IFN-β stimulation in the presence or absence of CHX, as described in (C). ISGF3 DNA-binding activity was normalized to IFN-β stimulation at 20 min.

Following IFN-β stimulation, phosphorylated STAT1 and STAT2 accumulated in both the cytoplasmic and nuclear fractions, consistent with sustained pathway activation and formation of the potentiated ISGF3 response. In contrast, IFN-λ3 stimulation resulted in a transient decrease in phosphorylated STAT1 and STAT2 during the same period in which nuclear ISGF3 DNA-binding activity entered the trough phase. Importantly, this decrease was apparent in the cytoplasmic fraction as well as the nuclear fraction. Thus, the IFN-λ3-specific trough in nuclear ISGF3 activity could not be explained solely by altered retention of activated STAT proteins.

These observations suggested that the mechanism responsible for the IFN-λ3 trough acts upstream of nuclear ISGF3 formation by transiently limiting STAT1 and STAT2 phosphorylation. We therefore considered whether differential engagement of a negative regulator of the JAK-STAT pathway could account for the distinct IFN-β and IFN-λ3 signaling trajectories.

To identify a potential negative regulator, we first examined whether the distinct IFN-β and IFN-λ3 signaling trajectories could be explained by differential induction of known negative regulators of the JAK-STAT signaling pathway (9, 10). We measured the temporal expression of SOCS1, SOCS3, and USP18 mRNA following stimulation with IFN-β or IFN-λ3 (Fig. 4B). SOCS1 was induced by both IFN-β and IFN-λ3 with similar induction during the first hour of stimulation. After one hour, however, IFN-β induced substantially higher SOCS1 expression than IFN-λ3. SOCS3 expression was induced by IFN-β but was not induced following IFN-λ3 stimulation. USP18 was induced by both IFN-β and IFN-λ3. Thus, none of the known negative regulators examined showed an IFN-λ3-specific expression pattern that could readily account for the transient inactivation of the pathway following IFN-λ3 stimulation.

### Tempered activation of ISGF3 during IFNλ3 stimulation is due to a latent negative regulator

Because transcriptional induction of these known negative regulators did not account for the IFN-λ3 signaling trajectory, we next sought to characterize the regulatory activity responsible for pathway inactivation, independent of the regulator’s identity. We reasoned that inhibiting protein synthesis would eliminate the contribution of newly synthesized positive and negative feedback regulators and thereby reveal the intrinsic stability of pathway activation following the two IFN stimuli. MLE-12 cells were therefore stimulated with IFN-λ3 in the presence or absence of the protein synthesis inhibitor cycloheximide (CHX), and nuclear ISGF3 DNA-binding activity and cytoplasmic phosphorylated STAT1 were measured over time (Fig. 4C).

In the absence of CHX, IFN-λ3 produced the characteristic ISGF3 response consisting of an initial activation followed by the trough and subsequent reactivation. In the presence of CHX, IFN-λ3 was still able to induce robust ISGF3 activity during the initial activation phase. ISGF3 activity initially exceeded that observed in the vehicle-treated condition, but after the first hour, it rapidly declined and ultimately returned to levels observed in unstimulated cells. Cytoplasmic pSTAT1 showed a similar response. Initial pSTAT1 activation was comparable in the presence and absence of CHX, followed by rapid inactivation and an eventual return to basal levels when protein synthesis was inhibited. Thus, blocking protein synthesis and preventing pathway reactivation by the positive feedback loops revealed rapid inactivation that occurs at the same time as the trough of the characteristic IFN-λ3 ISGF3 response.

We next performed the same experiment following IFN-β stimulation (Fig. 4D). As expected, IFN-β stimulation in the absence of CHX resulted in initial ISGF3 activation followed by the potentiated phase between approximately two and four hours. Consistent with our previous findings, inhibition of protein synthesis prevented this potentiated phase. However, unlike the IFN-λ3 response, ISGF3 activity did not rapidly decline following IFN-β stimulation in the presence of CHX. Instead, following the initial activation phase, ISGF3 activity remained relatively constant over the remainder of the measured time course. Cytoplasmic pSTAT1 similarly remained active in the presence of CHX.

Together, these results reveal a fundamental difference in the stability of JAK-STAT pathway activation following stimulation with IFN-β and IFN-λ3. In the absence of newly synthesized feedback regulators, IFN-β maintains ISGF3 and STAT1 activation, whereas IFN-λ3 signaling decays rapidly. Because inhibition of protein synthesis eliminates newly synthesized regulatory proteins yet does not prevent the rapid loss of IFN-λ3-induced STAT1 phosphorylation and ISGF3 activity, these results suggest that the IFN-λ3 pathway is subject to a pre-existing negative regulatory activity sufficient to drive pathway inactivation. In contrast, sustained IFN-β signaling appears resistant to this activity under the same conditions.

### The negative regulatory activity acts upstream of ISGF3 complex formation

The rapid loss of IFN-λ3-induced STAT phosphorylation and ISGF3 activity in the absence of protein synthesis suggested that a pre-existing regulatory activity limits IFN-λ3 signaling. We next sought to determine whether this regulatory activity acts directly on the ISGF3 complex or at an upstream step in the signaling pathway. Because IFN-β and IFN-λ3 activate the same ISGF3 transcription factor through distinct receptor complexes, we reasoned that simultaneous stimulation with both IFNs could distinguish between these possibilities. If the negative regulatory activity produces the IFN-λ3 trough by acting directly on ISGF3, then ISGF3 complexes generated during co-stimulation should remain susceptible to this regulation. Alternatively, if the regulatory activity acts upstream of ISGF3 formation in an IFN-λ3-specific manner, signaling through the IFN-β receptor should still be capable of generating the potentiated ISGF3 response during co-stimulation.

To test this, MLE-12 lung epithelial cells were stimulated with IFN-β, IFN-λ3, or both IFN-β and IFN-λ3 simultaneously, and nuclear ISGF3 DNA-binding activity was measured over time (Fig. 5A). Consistent with our previous experiments, IFN-β and IFN-λ3 produced similar levels of ISGF3 activity during the initial activation phase but subsequently generated distinct trajectories. IFN-β induced the potentiated phase of ISGF3 activity, whereas IFN-λ3 entered the characteristic trough. Interestingly, simultaneous stimulation with IFN-β and IFN-λ3 produced an ISGF3 trajectory like that observed with IFN-β alone. Co-stimulation did not substantially increase the initial ISGF3 response relative to either individual stimulus, but the response subsequently underwent potentiation rather than the trough observed with IFN-λ3.

**Figure 5.**
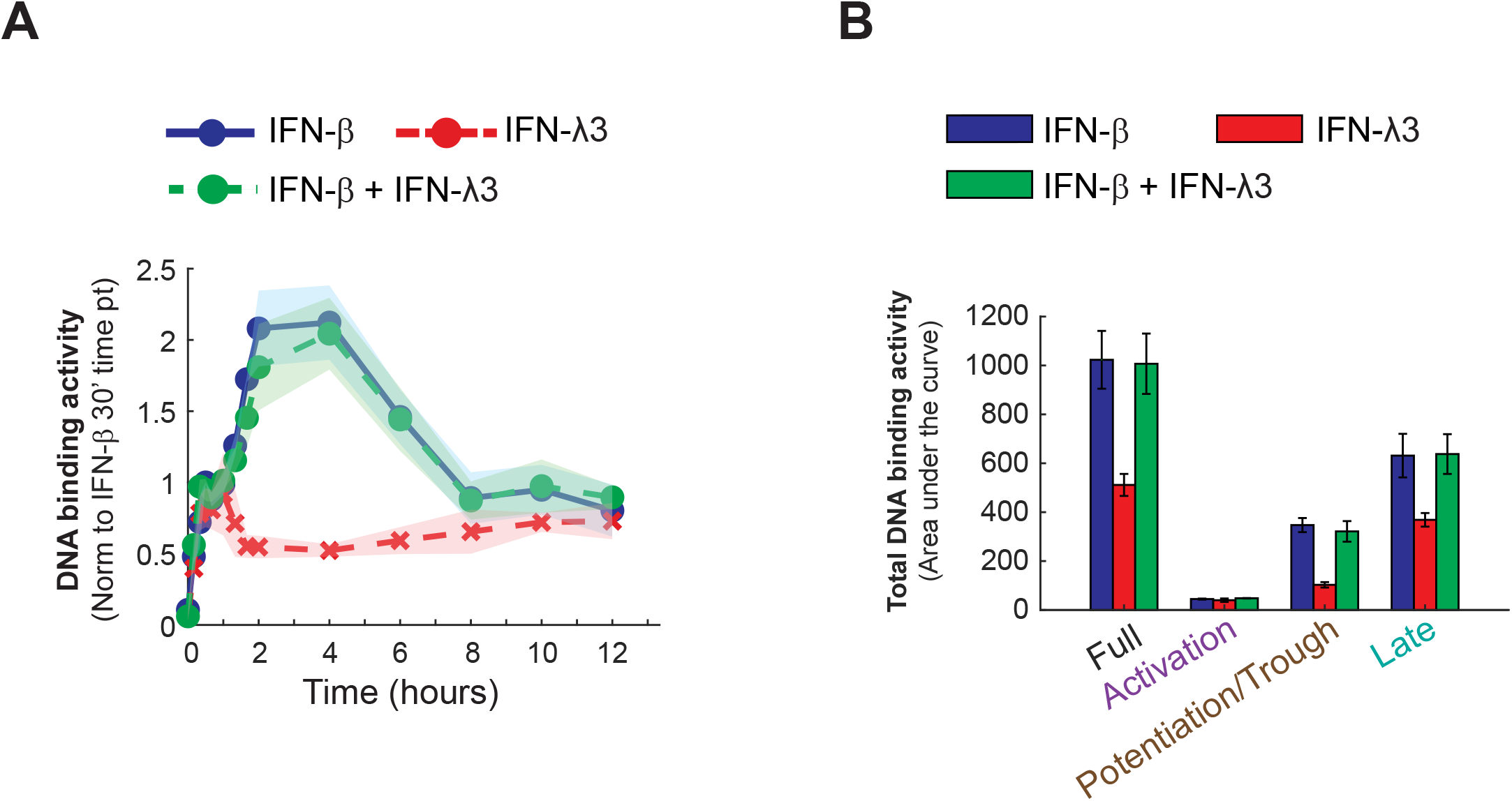
The negative regulator acting on the JAK-STAT pathway is not shared between the type I and type III IFN. A. ISGF3 DNA-binding activity following stimulation with 10 U/mL IFN-β (solid blue), 100 ng/mL IFN-λ3 (dashed red), or co-stimulation with 10 U/mL IFN-β and 100 ng/mL IFN-λ3 (dashed green). DNA-binding activity was measured by EMSA and normalized to IFN-β stimulation at 30 min. Shaded areas represent SEM from three independent experiments (n = 3). B. ISGF3 DNA-binding activity determined by AUC for the full time course and the activation, potentiation/trough, and late phases following stimulation as described in (A). Bars represent ± SEM from 3 independent experiments (n = 3).

To quantify these differences across the signaling trajectory, we calculated the area under the curves for the full ISGF3 activity response and for the individual activation and potentiation/trough phases (Fig. 5B). During the activation phase, integrated ISGF3 activity was similar among IFN-β, IFN-λ3, and co-stimulation conditions. In contrast, substantial differences emerged during the potentiation/trough phase. IFN-λ3 produced reduced integrated ISGF3 activity during this interval, whereas IFN-β and IFN-β/IFN-λ3 co-stimulation produced similarly elevated ISGF3 activity.

These results suggest that the pre-existing regulatory activity responsible for the IFN-λ3 trough does not act directly on ISGF3 complexes once they are formed. If IFN-λ3 activated a regulator that broadly inhibited ISGF3 itself, the ISGF3 response generated during co-stimulation would also be expected to undergo attenuation. Instead, IFN-β signaling maintained the potentiated ISGF3 response despite the simultaneous presence of IFN-λ3. Together, these results are consistent with a model in which the negative regulatory activity acts upstream of ISGF3 complex formation at an IFN-λ3-specific signaling step, rather than through a mechanism that indiscriminately inhibits ISGF3 activity.

### Pathway priming increases IFN-λ3-induced ISGF3 activation above the inflammatory gene activation threshold albeit transiently

Our results suggested that IFN-λ3 signaling is constrained by a negative regulatory activity independent of protein synthesis that acts upstream of ISGF3 formation. We next asked whether increasing the abundance of signaling pathway components prior to IFN-λ3 stimulation could generate sufficient ISGF3 activity to overcome this regulation and exceed the activation threshold associated with inflammatory gene induction. We reasoned that prior IFN exposure could increase the abundance of ISGF3 components, including STAT1, STAT2, and IRF9, thereby establishing a signaling state that could produce a more robust response to subsequent IFN-λ3 stimulation.

To test this, MLE-12 lung epithelial cells were primed with IFN-β, followed by neutralization of residual IFN-β and a rest period prior to secondary stimulation with IFN-λ3. Control conditions included cells stimulated with IFN-β or IFN-λ3 without prior priming, as well as primed cells that did not receive a secondary stimulation with IFN-λ3. This experimental design allowed us to distinguish residual signaling resulting from the initial IFN-β exposure from signaling newly induced by IFN-λ3 in the primed cellular state.

We first determined whether prior IFN exposure increased the abundance of ISGF3 components, including STAT1, STAT2, and IRF9. As anticipated, compared to the unprimed condition STAT1, STAT2, and IRF9 protein levels were increased in primed cells prior to secondary stimulation (Fig. 6A). Next, we assessed whether residual pathway activity remained following IFN-β priming and the subsequent rest period. Nuclear ISGF3 DNA-binding activity was low in primed cells prior to secondary IFN-λ3 stimulation, indicating that the priming protocol did not result in substantial persistent formation of transcriptionally active ISGF3 complexes (Fig. 6B). Similarly, cytoplasmic phosphorylated STAT1 and STAT2 were low following the rest period, although a modest elevation in phosphorylated STAT2 was observed in primed compared with unprimed cells. Thus, the cellular state established by IFN-β priming could be examined without substantial residual activation of the JAK-STAT pathway from the initial stimulus.

**Figure 6.**
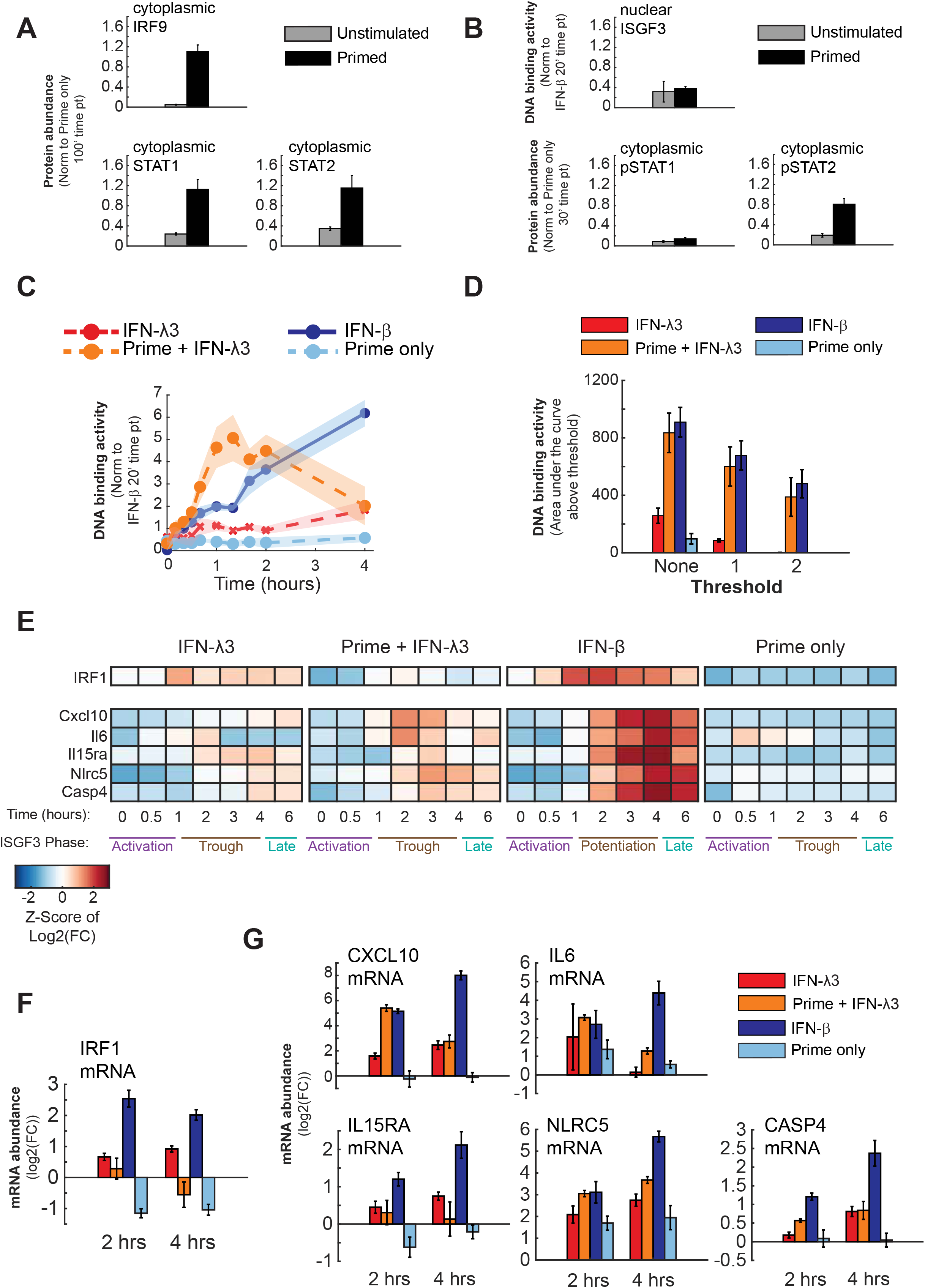
Although priming the signaling pathway increases initial ISGF3 activation, rapid pathway inactivation still leads to limited/no inflammatory gene activation (?) A. Cytoplasmic IRF9, STAT1, and STAT2 abundance in unstimulated cells (gray bars) and cells primed with 10 U/mL IFN-β for 16 h (black bars). Protein abundance was measured by western blotting and normalized to 100 min time point. Bars represent ± SEM from 3 independent experiments (n = 3). B. ISGF3 DNA-binding activity and cytoplasmic pSTAT1 and pSTAT2 abundance in unstimulated and IFN-β-primed cells, as described in (A). ISGF3 DNA-binding activity was measured by EMSA and normalized to the 20 min IFN-β stimulation. Protein abundance was quantified by western blotting and normalized to IFN-β stimulation at 30 min. Bars represent mean ± SEM from three independent experiments (n = 3). C. ISGF3 activity following stimulation with 10 U/mL IFN-β (solid blue) or 100 ng/mL IFN-λ3 (dashed red), with or without IFN-β priming. Following priming, IFN-β was neutralized with an IFN-β-neutralizing antibody (Nab; 60 NU/mL), and cells were rested before stimulation with IFN-λ3. Primed cells without subsequent cytokine stimulation were included as a control. DNA-binding activity was measured by EMSA and normalized to IFN-β stimulation at 20 min. Shaded areas represent SEM from at least 3 independent experiments (n = 3). D. AUC quantification of ISGF3 DNA-binding activity shown in (C) over the full time course above indicated thresholds. Bars represent mean ± SEM from three independent experiments. E. Heatmap of IRF1 and selected pro-inflammatory gene expression following IFN-λ3 stimulation, IFN-β priming followed by IFN-λ3 stimulation, IFN-β stimulation, or priming alone, as described in (C). mRNA abundance was measured by RT-qPCR and normalized as described in Figure 2A. Log2 fold-change values were Z-score normalized for each gene for visualization. The corresponding ISGF3 signaling phases are indicated below the heatmaps. F. IRF1 mRNA expression at 2 and 4 h for the conditions described in (E). Bars represent ± SEM from 3 independent experiments (n = 3). G. CXCL10, IL6, IL15RA, NLRC5, and CASP4 mRNA expression at 2 and 4 h for the conditions shown in (E). Bars represent ± SEM from 3 independent experiments (n = 3).

Following secondary stimulation with IFN-λ3, primed cells increased the initial amplitude of IFN-λ3-induced ISGF3 activity but did not eliminate the subsequent attenuation of the pathway (Fig. 6C). ISGF3 activity began to plateau at approximately one hour and decreased after two hours. Priming cells before the IFN-λ3 stimulation was sufficient to activate ISGF3 DNA-binding activity above the threshold previously associated with inflammatory gene activation (Fig. 6D). This result was consistent with the prediction that increasing pathway competence prior to stimulation could amplify the subsequent IFN-λ3 response.

### Pathway priming enables IFN-λ3 to transiently induce inflammatory gene expression

We next determined whether the increased ISGF3 activity generated by IFN-β priming altered downstream transcriptional responses following IFN-λ3 stimulation. We first examined IRF1, which has been implicated in the differential inflammatory response to type I and type III IFNs. Interestingly, following IFN-β priming, neutralization, and the 10-hour rest period, IRF1 gene expression was lower than in cells that had not undergone priming (Fig. 6E). Thus, the primed cellular state was not characterized by elevated basal IRF1 gene expression.

Secondary stimulation of primed cells with IFN-λ3 induced IRF1 gene expression above the primed baseline. However, the amount of IRF1 induced remained lower than that observed following IFN-β stimulation and lower than that observed following IFN-λ3 stimulation in unprimed cells. Thus, although IFN-β priming increased the subsequent ISGF3 DNA binding activity by IFN-λ3, it did not similarly increase IRF1 mRNA expression. This provided an opportunity to determine whether the increased ISGF3 response in primed cells was nevertheless associated with induction of inflammatory genes.

We therefore measured the expression of CXCL10, IL6, IL15RA, NLRC5, and CASP4 following IFN-β, IFN-λ3, or IFN-λ3 stimulation of IFN-β-primed cells (Fig. 6E). Despite the reduced IRF1 expression in the primed condition, secondary IFN-λ3 stimulation induced expression of these inflammatory genes. At two hours, the primed plus IFN-λ3 condition produced levels of CXCL10, IL6, and NLRC5 expression that were similar to those induced by IFN-β (Fig. 7G). However, this similarity was transient. By four hours, IFN-β induced greater expression of these genes than the primed plus IFN-λ3 condition. The primed IFN-λ3 response therefore acquired the capacity to induce an inflammatory gene expression program but did not reproduce the magnitude or duration of the response induced by IFN-β. The transient nature of this transcriptional response corresponded with the ISGF3 signaling trajectory (Fig. 7C). Thus, the early increase in ISGF3 activity in primed cells was associated with inflammatory gene induction at two hours, whereas subsequent attenuation of ISGF3 activity was associated with divergence of the inflammatory transcriptional response from that induced by IFN-β.

Together, these results suggest that changing the cellular state prior to IFN-λ3 stimulation can alter the transcriptional response to the cytokine by changing ISGF3 temporal dynamics. Increasing pathway responsiveness through IFN-β priming enabled IFN-λ3 to generate sufficient ISGF3 activity above the gene activation threshold to induce inflammatory genes that are otherwise weakly induced. Importantly, this occurred despite reduced IRF1 gene expression, suggesting that the enhanced inflammatory response in the primed condition is associated with increased ISGF3 activity rather than enhanced IRF1 induction. However, priming did not prevent the subsequent attenuation of IFN-λ3 signaling, resulting in a less robust and shorter-lived inflammatory response than that induced by IFN-β. These results further support a model in which both the magnitude and duration of ISGF3 activity above a gene specific activation threshold contribute to the stimulus-specific regulation of inflammatory gene expression.

## DISCUSSION

Here, we identify a temporal signaling mechanism that contributes to the distinction between type I and type III IFN responses. Our results suggest that the inflammatory output of the IFN/JAK-STAT pathway is not determined solely by whether ISGF3 or IRF1 is activated. Instead, inflammatory genes are associated with a more stringent signaling requirement in which ISGF3 activity must remain above an activation threshold during a critical portion of the response. IFN-β satisfies this requirement through a potentiated phase of ISGF3 activity, whereas IFN-λ3 undergoes a trough that limits both the magnitude and duration of activity above the threshold. This provides a mechanistic framework for understanding how two cytokine families that converge on the same transcription factor can retain a shared antiviral program while differing in their capacity to induce inflammation.

This work extends previous models of type I and type III IFN specificity. Forero et al. showed that low IFNLR1 abundance limits STAT1 activation and IRF1 induction and proposed an IRF1-dependent threshold for inflammatory gene activation (5)(Forero et al., 2019). Our data support the importance of IRF1 but indicate that IRF1 induction alone does not fully account for inflammatory selectivity. In lung epithelial cells, IFN-λ3 induced IRF1 during the first hour, but expression was rapidly truncated. More importantly, IFN-β priming enabled subsequent IFN-λ3 stimulation to induce CXCL10, IL6, IL15RA, NLRC5, and CASP4 even though IRF1 protein abundance remained lower than in the IFN-β condition. The inflammatory response was strongest at early times and became less robust as ISGF3 activity declined. Thus, IRF1 and ISGF3 should not be considered independent binary switches. Recent work showing that IRF1 can cooperate with ISGF3 to establish innate immune enhancers provides additional support for considering their activities together and over time (11). Consistent with our findings, time-dependent recruitment of ISGF3, GAF, and IRF1 to distinct and composite regulatory elements has been shown to shape IFN-induced transcriptional programs, further supporting a model in which mechanisms that control the magnitude and duration of IFN-activated transcription factors can determine which gene programs are engaged (12).

Our findings also help reconcile apparently different conclusions regarding receptor abundance. Receptor abundance clearly affects signaling amplitude and can determine whether a pathway reaches a transcriptional threshold. However, Pervolaraki et al. found that receptor abundance did not explain the distinct kinetics of type I and type III IFN responses (6). Consistent with this distinction, increasing the IFN-λ3 dose in our experiments increased pathway activation at lower concentrations but did not remove the trough once the response approached saturation. We therefore propose that receptor abundance sets an upper signaling capacity, whereas receptor-proximal and intracellular regulatory mechanisms determine the trajectory produced within that capacity. This distinction between how much signaling can occur and how signaling changes over time may be particularly important for tunable genes with higher activation requirements. Prior work has shown that IFN-dependent accumulation of ISGF3 components contributes to positive feedback that sustains IFN signaling, with IRF9 and STAT2 playing important roles in ISGF3 potentiation (8) However, our findings suggest that insufficient engagement of this positive feedback does not readily explain the IFN-λ3 trough. STAT1, STAT2, and IRF9 mRNA and protein remained induced following IFN-λ3 stimulation, and the cytoplasmic availability of these ISGF3 components was not reduced during the trough phase. Together, these findings suggest that increasing the availability of positive-feedback components is insufficient to prevent the temporal reduction of IFN-λ3-induced ISGF3 activity and point toward an additional regulatory mechanism controlling this response.

The cycloheximide and IFN co-stimulation experiments further localize the mechanism that constrains the IFN-λ3 trajectory. Inhibition of protein synthesis prevented the IFN-β potentiated phase, consistent with its requirement for induced positive feedback, while IFN-β signaling remained comparatively stable. In contrast, IFN-λ3-induced pSTAT1 and ISGF3 activity rapidly decayed when protein synthesis was inhibited. Because this decay occurred without synthesis of new feedback regulators, these data point to a negative regulatory activity independent of protein synthesis. The corresponding decrease in cytoplasmic phosphorylated STAT1 and STAT2 places this activity upstream of nuclear ISGF3 accumulation. Moreover, simultaneous IFN-β and IFN-λ3 stimulation produced an IFN-β-like, ISGF3-potentiated trajectory, arguing against indiscriminate inhibition of ISGF3 after the two pathways converge. Together, these observations are most consistent with regulation at an IFN-λ3-selective, receptor-proximal step upstream of STAT phosphorylation and ISGF3 formation.

Several known regulators of IFN-λ signaling provide potential mechanisms for this activity. FBXO45 directly associates with the intracellular domain of IFNLR1 and promotes its ubiquitination and proteasomal degradation in airway epithelial cells; FBXO45 depletion increases IFN-λ-induced STAT1 phosphorylation and ISG expression, whereas its overexpression selectively suppresses IFN-λ signaling (13). Interestingly, that study identified FBXO45 using two independent proximity-labeling approaches and mapped its interaction to the IFNLR1 intracellular domain, providing evidence that receptor-associated protein turnover can regulate IFN-λ signaling upstream of STAT1. SOCS1 also negatively regulates IFN-λ signaling by acting at the receptor-JAK complex, and SOCS1 deficiency increases IFN-λ-induced ISG expression in cells and responsive tissues in vivo (9). However, in our study SOCS1 transcriptional induction was not specific to IFN-λ3 making this mechanism not an obvious explanation for the rapid attenuation we observe when new protein synthesis is blocked.

Protein tyrosine phosphatases provide another potential mechanism for rapid regulation by proteins already present when IFN-λ3 stimulation begins. SHP2/PTPN11 can negatively regulate IFN-induced JAK-STAT signaling, as loss of functional SHP2 increases STAT activation and STAT1 phosphorylation following IFN stimulation (14). Although direct recruitment of SHP2 to IFNLR1 and selective regulation of IFN-λ signaling have not been established, its phosphotyrosine-recognizing SH2 domains and ability to constrain IFN-induced STAT activation make receptor- or kinase-associated phosphatase activity a plausible mechanism for the rapid signal attenuation observed here.

Thus, the molecular identity of the activity responsible for the IFN-λ3 trajectory remains unresolved. Our findings suggest that IFN-λ3 signaling may be constrained by a pre-existing receptor-proximal regulator that transiently reduces signaling through the IFNLR complex and thereby limits the magnitude and duration of STAT phosphorylation and ISGF3 activity. Such regulation could operate alongside established mechanisms controlling IFNLR1 abundance and inducible negative feedback, providing an additional layer through which epithelial cells restrict the time that IFN-λ3-induced ISGF3 activity remains above the threshold required for inflammatory gene activation.

The priming experiments demonstrate that the IFN-λ3 response is not fixed. Prior IFN-β exposure altered the pathway state, such that a subsequent IFN-λ3 stimulus elicited a larger initial ISGF3 response and transient induction of inflammatory genes. However, priming did not eliminate the later attenuation of IFN-λ3 signaling. This separation is informative: increasing pathway competence can move IFN-λ3 across the inflammatory activation threshold, but the IFN-λ3-specific negative regulatory activity still constrains how long the response remains there. Prior studies have shown that IFN exposure can alter subsequent cytokine responsiveness through feedback and refractory states, and viral infection naturally generates temporally changing and overlapping type I and type III IFN environments. Our results provide a mechanistic basis for how such exposure history could change the functional meaning of a later IFN signal without changing the identity of the ligand itself. This temporal framework may also help explain the division of labor between antiviral protection and inflammation observed in vivo. During respiratory infection, type III IFNs can provide early barrier-focused antiviral protection, whereas type I IFNs are more strongly associated with later inflammatory responses (2). Our data suggest that cells may distinguish these inputs not simply by measuring ligand or receptor abundance but by decoding trajectory features of the shared JAK-STAT pathway. In this model, antiviral genes with lower or more permissive activation requirements remain responsive to both IFN types, whereas inflammatory genes act as tunable outputs that require sufficiently strong and sustained ISGF3 activity.

There are several limitations to the present study. The experiments were performed primarily in MLE-12 lung epithelial cells, and the molecular identity of the pre-existing IFN-λ3-specific regulatory activity remains unresolved. The threshold analysis also does not imply that all inflammatory genes share a single quantitative threshold; individual genes are likely to integrate ISGF3 with additional transcription factors, chromatin state, and gene-specific regulatory mechanisms. IRF1 remains an important contributor to inflammatory gene regulation, and our results do not argue that ISGF3 dynamics replace IRF1-dependent control. Rather, they indicate that the duration and amplitude of ISGF3 activity provide an additional regulatory dimension that can determine whether the signaling environment is sufficient for robust inflammatory transcription. Future studies aimed at identifying the regulatory mechanism responsible for the IFN-λ3 transient reduction in signaling will be important for testing our proposed model. Defining this mechanism may also have important implications for how type III IFN responses are therapeutically modulated. Understanding how IFN-λ3 signaling is temporally regulated could therefore provide opportunities to selectively modify the duration or strength of JAK-STAT signaling without broadly disrupting the shared signaling pathway. More broadly, understanding how closely related cytokines generate distinct outputs through shared signaling components may provide a framework for developing therapies that target signaling dynamics rather than the complete activation or inhibition of pathways.

## METHODS

### Key resources table

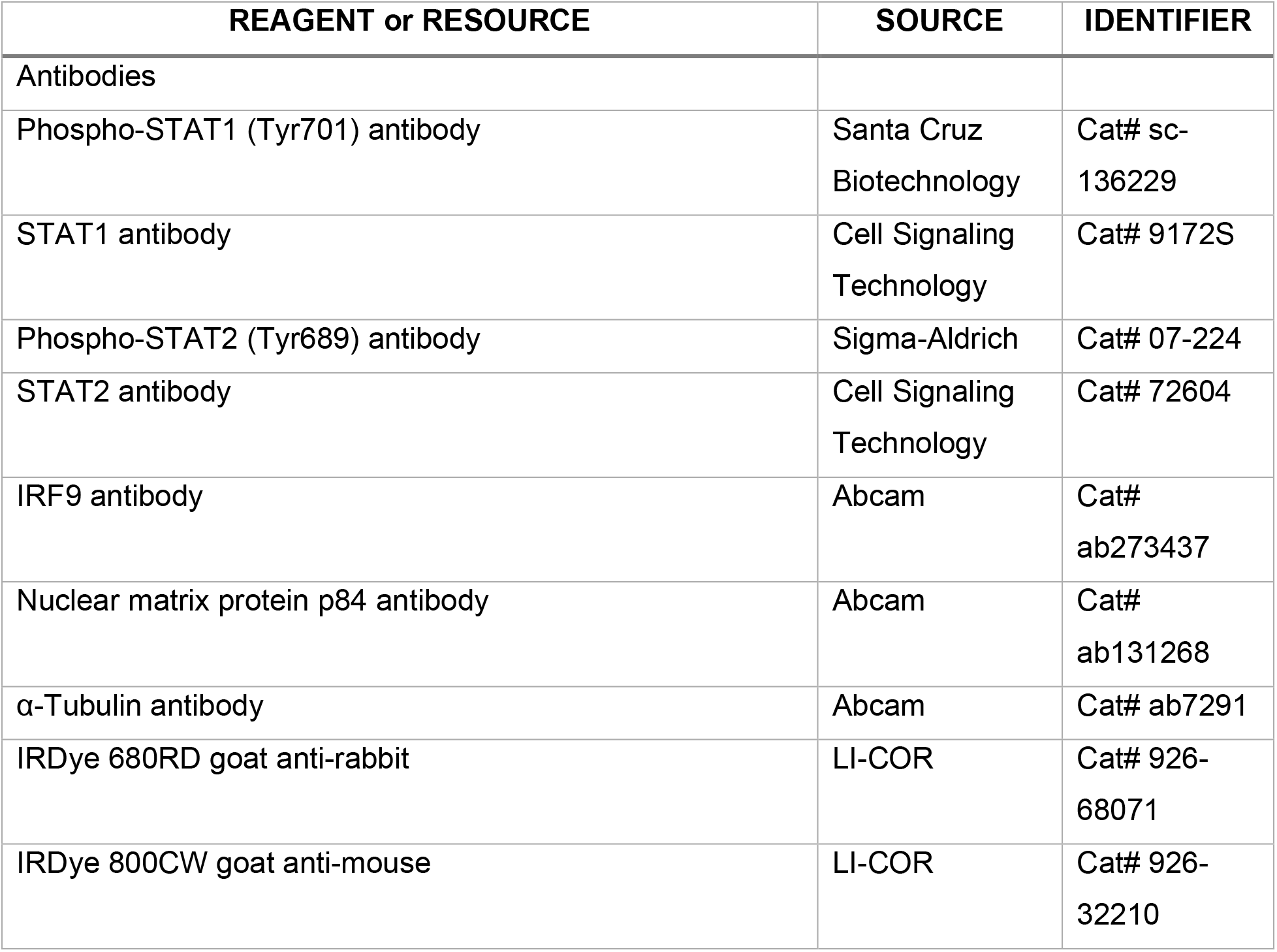
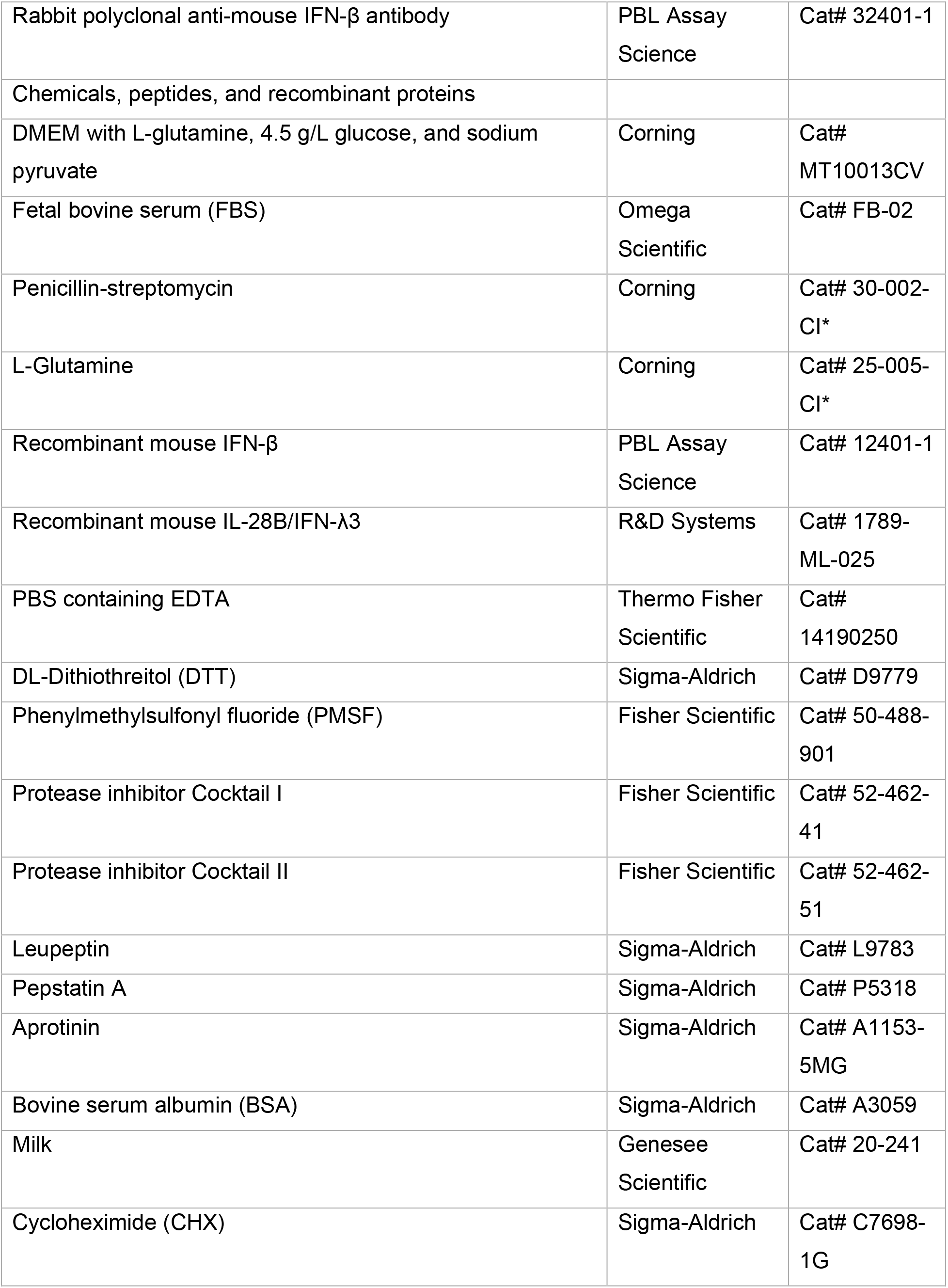
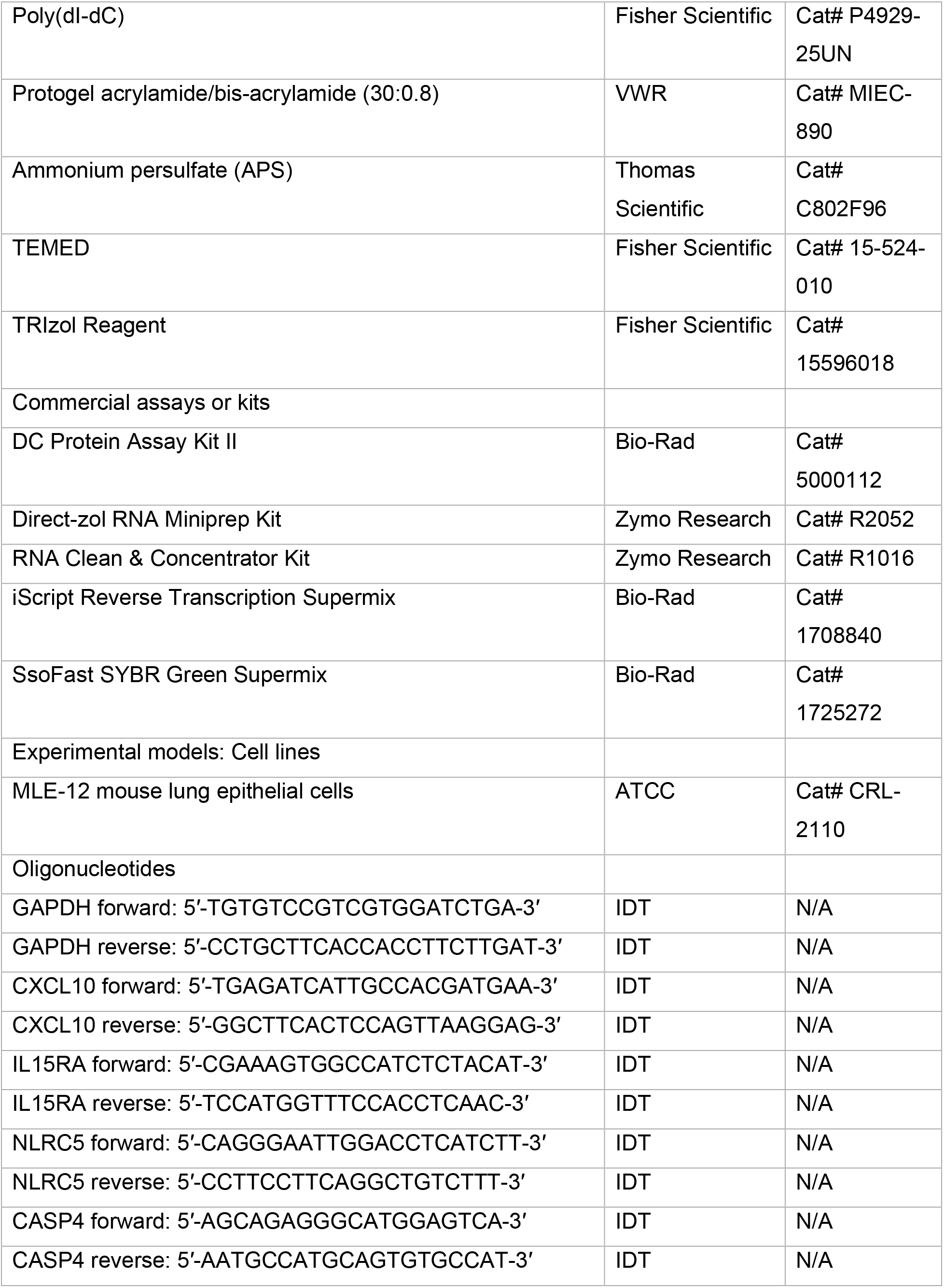
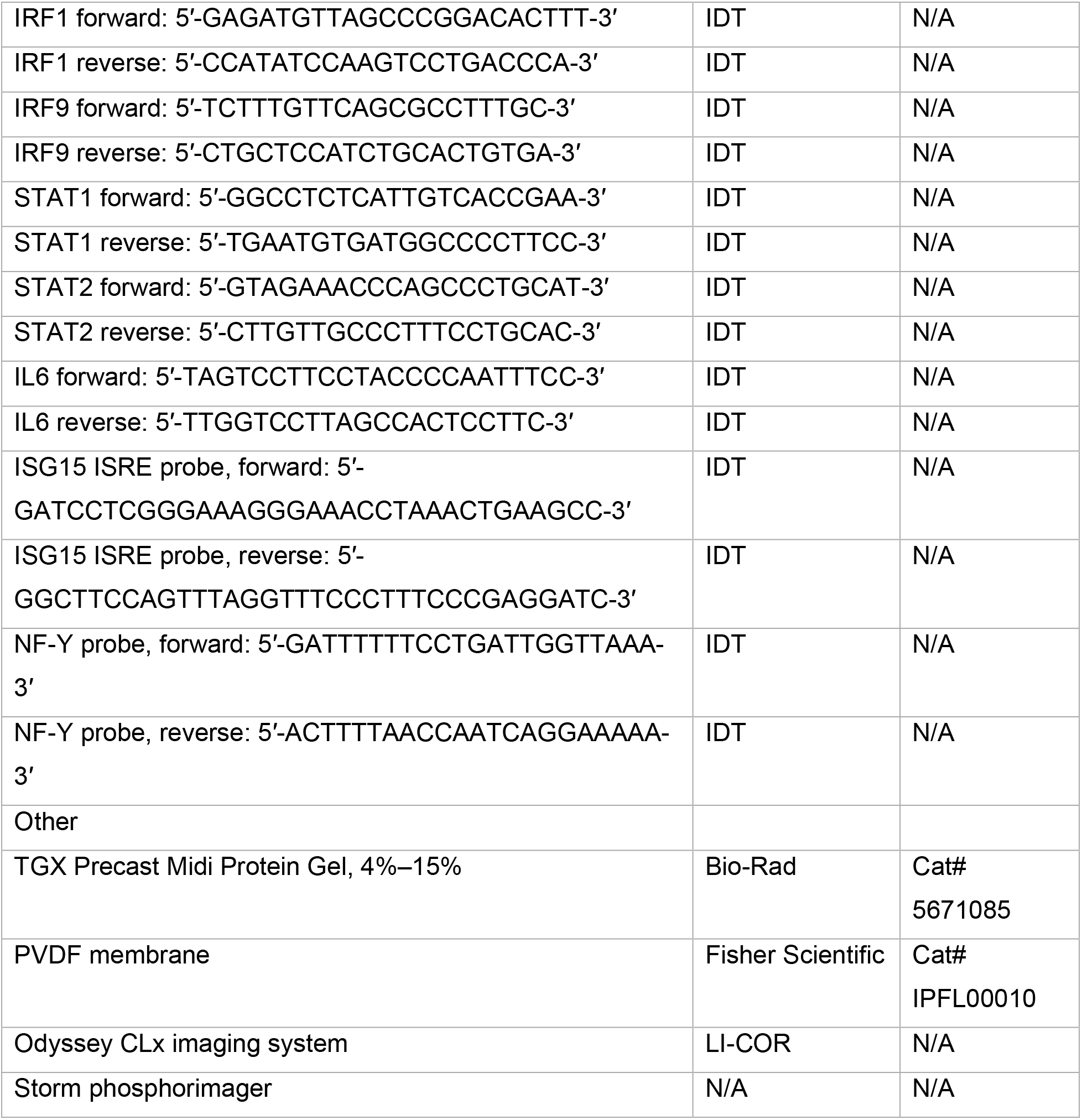

### Cell culture and cytokine stimulation

MLE-12 mouse lung epithelial cells were cultured in DMEM containing L-glutamine, 4.5 g/L glucose, and sodium pyruvate, supplemented with 10% fetal bovine serum (FBS), 100 IU/mL penicillin, 100 μg/mL streptomycin, and 2 mM L-glutamine. Cells were maintained at 37°C in a humidified incubator with 5% CO₂.

For cytokine stimulation, cells were treated with recombinant mouse IFN-β (0.1–100 U/mL) or IL-28B/IFN-λ3 (0.1–100 ng/mL), as indicated. For priming experiments, cells were treated with 10 U/mL IFN-β for 16 h, followed by the addition of a neutralizing anti-mouse IFN-β antibody. Cells were rested for 10 h and subsequently stimulated with 100 ng/mL IL-28B/IFN-λ3.

For cycloheximide (CHX) experiments, cells were stimulated with 100 ng/mL IFN-λ3 or 10 U/mL IFN-β in the presence or absence of 20 ug/mL CHX. CHX and either IFN-λ3 or IFN-β were added simultaneously without CHX pretreatment. Samples were collected at 0, 10, 30, 60, 80, and 100 min and 2, 4, 6, 8, 10, and 12 h after stimulation.

For co-stimulation experiments, cells were treated with 100 ng/mL IFN-λ3, 10 U/mL IFN-β, or both cytokines simultaneously. Samples were collected at 0, 10, 20, 30, 40, 60, 80, and 100 min, and at 2, 4, 6, 8, 10, and 12 h after stimulation.

### Cytoplasmic and nuclear protein extraction

Following cytokine stimulation, cells were washed with ice-cold PBS containing 1 mM EDTA and collected by centrifugation. Cell pellets were resuspended in 80 μL cytoplasmic extraction (CE) buffer supplemented with 1 mM DTT, 1 mM PMSF, protease inhibitor Cocktails I and II (10 μL/mL each), 4 μg/mL leupeptin, 0.001 mM pepstatin A, and 0.01 TIU/mL aprotinin. Samples were incubated on ice for 2 min, vortexed for 30 s, and centrifuged for 30 s, and the supernatant was collected as the cytoplasmic fraction. Remaining pellet was washed twice with 80 μL CE buffer and resuspended in 20–30 μL nuclear extraction (NE) buffer containing the same inhibitors. Samples were subjected to three freeze–thaw cycles (5 min on dry ice followed by 30 s at 37°C) and centrifuged at maximum speed for 5 min at 4°C. The supernatant was collected as the nuclear fraction.

### Western blotting

Protein concentrations were determined using the DC Protein Assay Kit II. Equal amounts of protein were loaded on 4%–15% gradient SDS-PAGE gels and transferred to PVDF membranes at 100 V for 1 h. Membranes were blocked for 1 h at room temperature in 5% BSA or 5% milk in TBST and incubated overnight at 4°C with primary antibodies against phospho-STAT1 (Tyr701; 1:1,000), STAT1 (1:1,000), phospho-STAT2 (Tyr689; 1:200), STAT2 (1:500), IRF9 (1:1,000), p84 (1:5,000), or α-tubulin (1:5,000). Membranes were washed with TBST containing 0.5% Tween-20, incubated with IRDye secondary antibodies for 1 h at room temperature, washed, and imaged using an Odyssey CLx imaging system (LI-COR)

### Electrophoretic mobility shift assay

Electrophoretic mobility shift assays (EMSAs) were performed as previously described (Hoffmann et al., 2002; Werner et al., 2005; Wilder et al., 2023). Equal amounts of nuclear protein were incubated for 30 min with 0.01282 pmol of ^32P-labeled double-stranded oligonucleotide probe in binding buffer containing 10 mM Tris-HCl (pH 7.5), 50 mM NaCl, 10% glycerol, 1% IGEPAL-630, 1 mM EDTA (pH 8.0), and 0.1 mg/mL poly(dI-dC). Probes contained either an ISRE sequence derived from the ISG15 locus or an NF-Y binding sequence; oligonucleotide sequences are provided in the Key Resources Table. Samples were loaded onto a 5% polyacrylamide gels (30:0.8 acrylamide:bis-acrylamide) containing 5% glycerol in 1× TGE buffer (25 mM Tris, 190 mM glycine, and 1 mM EDTA, pH 8.0) at 200 V. Following electrophoresis, gels were dried, and radiolabeled protein-DNA complexes were visualized by phosphorimaging using a Storm phosphorimager (UCSF Parnassus Stem Cell Core, Blelloch Lab).

### RNA isolation, cDNA synthesis, and qPCR

RNA was harvested from cells lysed in TRIzol using the Direct-zol RNA Miniprep Kit and further purified using the RNA Clean & Concentrator Kit as per the manufacturer’s instructions. cDNA was synthesized from 100 ng total RNA using iScript Reverse Transcription Supermix and diluted 1:8 for quantitative PCR (qPCR) using SsoFast SYBR Green Supermix. Relative gene expression was calculated using the 2^−ΔΔCt method, with *Gapdh* as the reference gene. Gene-specific primers are listed in the Key Resources Table.

## CONFLICT OF INTEREST

The authors declare no conflicts of interest.

## FUNDING INFORMATION

This research was funded by the UCLA Chancellor’s Postdoctoral Fellowship Program and start-up funds from the University of California, San Francisco and Biohub, San Francisco, Investigator Award.

## SUPLEMENTAL FIGURES

**Figure S1. IFN-β and duration modulate ISGF3 signaling dynamics.**

A. ISGF3 DNA-binding activity following stimulation with IFN-β (10 U/mL, solid blue), low-dose IFN-β (1 U/mL, dashed light blue), or a 15-min pulse of 10 U/mL IFN-β (dashed purple). DNA-binding activity was measured by EMSA and normalized to 10 U/mL IFN-β stimulation at 20 min. Shaded areas represent SEM from at least three independent experiments (n = 3 - 10).

B. Schematic of area-under-the-curve (AUC) analysis used to quantify ISGF3 DNA-binding activity over the full time course (i) and the activation (ii), potentiation/trough (iii), and late (iv) phases.

C. AUC quantification of ISGF3 DNA-binding activity shown in (A) over the full time course and the activation, potentiation/trough, and late phases defined in (B). Bars represent mean ± SEM from at least three independent experiments (n = 3 - 10).

## Notes

### Competing Interest Statement

The authors have declared no competing interest.

## REFERENCES

1. Lazear HM, Schoggins JW, Diamond MS. Shared and Distinct Functions of Type I and Type III Interferons. Immunity. 2019;50(4):907–23. doi: 10.1016/j.immuni.2019.03.025. PubMed PMID: 30995506; PubMed Central PMCID: PMC6839410.

2. Galani IE, Triantafyllia V, Eleminiadou EE, Koltsida O, Stavropoulos A, Manioudaki M, et al. Interferon-lambda Mediates Non-redundant Front-Line Antiviral Protection against Influenza Virus Infection without Compromising Host Fitness. Immunity. 46. United States: 2017 Elsevier Inc; 2017. p. 875–90 e6.

3. Mesev EV, LeDesma RA, Ploss A. Decoding type I and III interferon signalling during viral infection. Nat Microbiol. 2019;4(6):914–24. Epub 20190401. doi: 10.1038/s41564-019-0421-x. PubMed PMID: 30936491; PubMed Central PMCID: PMC6554024.

4. Aaronson DS, Horvath CM. A road map for those who don’t know JAK-STAT. Science. 2002;296(5573):1653–5. doi: 10.1126/science.1071545. PubMed PMID: 12040185.

5. Forero A, Ozarkar S, Li H, Lee CH, Hemann EA, Nadjsombati MS, et al. Differential Activation of the Transcription Factor IRF1 Underlies the Distinct Immune Responses Elicited by Type I and Type III Interferons. Immunity. 2019;51(3):451–64.e6. Epub 20190827. doi: 10.1016/j.immuni.2019.07.007. PubMed PMID: 31471108; PubMed Central PMCID: PMC7447158.

6. Pervolaraki K, Rastgou Talemi S, Albrecht D, Bormann F, Bamford C, Mendoza JL, et al. Differential induction of interferon stimulated genes between type I and type III interferons is independent of interferon receptor abundance. PLoS Pathog. 2018;14(11):e1007420. Epub 20181128. doi: 10.1371/journal.ppat.1007420. PubMed PMID: 30485383; PubMed Central PMCID: PMC6287881.

7. Purvis JE, Lahav G. Encoding and decoding cellular information through signaling dynamics. Cell. 2013;152(5):945–56. doi: 10.1016/j.cell.2013.02.005. PubMed PMID: 23452846; PubMed Central PMCID: PMC3707615.

8. Wilder CL, Lefaudeux D, Mathenge R, Kishimoto K, Zuniga Munoz A, Nguyen MA, et al. A stimulus-contingent positive feedback loop enables IFN-β dose-dependent activation of pro-inflammatory genes. Mol Syst Biol. 2023:e11294. Epub 20230317. doi: 10.15252/msb.202211294. PubMed PMID: 36929731.

9. Blumer T, Coto-Llerena M, Duong FHT, Heim MH. SOCS1 is an inducible negative regulator of interferon λ (IFN-λ)-induced gene expression in vivo. J Biol Chem. 2017;292(43):17928–38. PubMed PMID: 28900038.

10. Kubo M, Hanada T, Yoshimura A. Suppressors of cytokine signaling and immunity. Nat Immunol. 4. United States 2003. p. 1169-76.

11. Chavez C, Lin K, Malveaux A, Gorin A, Brizuela S, Cheng QJ, et al. IRF1 cooperates with ISGF3 or GAF to form innate immune de novo enhancers in macrophages. Sci Signal. 2025;18(868):eado8860. Epub 20250107. doi: 10.1126/scisignal.ado8860. PubMed PMID: 39772531; PubMed Central PMCID: PMC12052582.

12. Sekrecka A, Kluzek K, Sekrecki M, Boroujeni ME, Hassani S, Yamauchi S, et al. Time-dependent recruitment of GAF, ISGF3 and IRF1 complexes shapes IFNα and IFNγ-activated transcriptional responses and explains mechanistic and functional overlap. Cell Mol Life Sci. 2023;80(7):187. Epub 20230622. doi: 10.1007/s00018-023-04830-8. PubMed PMID: 37347298; PubMed Central PMCID: PMC10287828.

13. Tsai M, Osman W, Adair J, ElMergawy R, Chafin L, Johns F, et al. The E3 ligase subunit FBXO45 binds the interferon-λ receptor and promotes its degradation during influenza virus infection. J Biol Chem. 2022;298(12):102698. Epub 20221113. doi: 10.1016/j.jbc.2022.102698. PubMed PMID: 36379255; PubMed Central PMCID: PMC9747586.

14. You M, Yu DH, Feng GS. Shp-2 tyrosine phosphatase functions as a negative regulator of the interferon-stimulated Jak/STAT pathway. Mol Cell Biol. 1999;19(3):2416–24. doi: 10.1128/MCB.19.3.2416. PubMed PMID: 10022928; PubMed Central PMCID: PMC84034.

