## Supplementary figures and images for "Type I and III interferons differentially encode inflammatory gene activation through threshold control of ISGF3 dynamics"

### Supplemental Figure 1

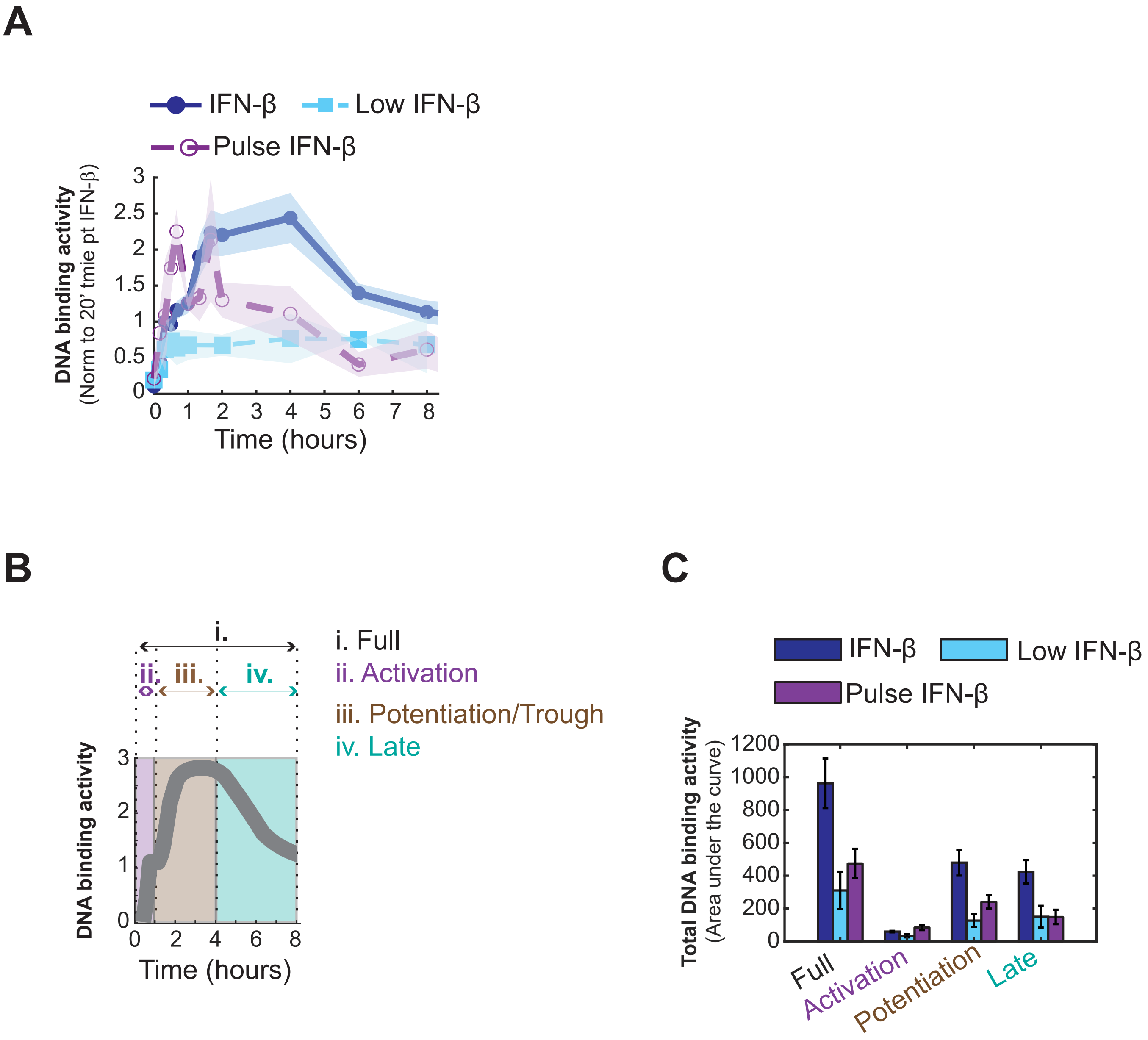
